# Rhythmic temporal coordination of neural activity prevents representational conflict during working memory

**DOI:** 10.1101/2022.12.02.518876

**Authors:** Miral Abdalaziz, Zach V. Redding, Ian C. Fiebelkorn

## Abstract

Selective attention^1^ is characterized by alternating states associated with either attentional sampling or attentional shifting, helping to avoid functional conflicts by isolating function-specific neural activity in time^2–5^. We hypothesized that such rhythmic temporal coordination might also help to avoid representational conflicts during working memory^6^. Multiple items can be simultaneously held in working memory, and these items can be represented by overlapping neural populations^7–9^. Traditional theories propose that short-term storage of to-be-remembered items occurs through persistent neural activity^10–12^, but when neurons are simultaneously representing multiple items, persistent activity creates a potential for representational conflicts. In comparison, more recent, ‘activity-silent’ theories of working memory propose that synaptic changes also contribute to the short-term storage of to-be-remembered items^13–16^. Transient bursts in neural activity^17^, rather than persistent activity, could serve to occasionally refresh these synaptic changes. Here, we used EEG and response times (RTs) to test whether rhythmic temporal coordination helps to isolate neural activity associated with different to-be-remembered items, which would help to avoid representational conflicts. Consistent with this hypothesis, we report that the relative strength of different item representations alternates over time as a function of frequency-specific phase. Although RTs were linked to theta (~6Hz) and beta (~25 Hz) phase during a memory delay, the relative strength of item representations only alternated as a function of beta phase. The present findings (i) are consistent with rhythmic temporal coordination being a general mechanism for avoiding either functional or representational conflicts during cognitive processes, and (ii) inform models describing the role of oscillatory dynamics in organizing working memory^13,18–21^.

## RESULTS

The rhythmic coordination of neural activity can help to isolate competing functions or information in time^22–27^. For example, previous studies have demonstrated that selective attention is characterized by rhythmically alternating states and associated fluctuations in perceptual sensitivity (at ~4-6 Hz)^3–5,28–37^. These alternating states help to temporally isolate potentially conflicting sensory (i.e., attention-related sampling) and motor (i.e., attention-related shifting) functions within the network that directs both attention-related changes in sensory processing and orienting movements (i.e., the ‘attention network’)^3^. Here, we tested whether rhythmic temporal coordination of neural activity is a more general mechanism for avoiding conflict during cognitive processes. Specifically, we tested whether such coordination helps to temporally isolate item-specific neural activity during working memory, which would help to avoid *representational conflicts*.

Working memory is the process through which behaviorally important information is temporarily stored and internally sampled^6,38^. Traditional theories of working memory have proposed that the short-term storage of a to-be-remembered item occurs through persistent neural activity^10^, where persistent neural activity is defined as increased spiking activity that spans an entire memory delay (occurring among neurons representing the to-be-remembered item). While there is clear evidence that some neurons demonstrate such persistent activity during working memory delays^10–12,39,40^, short-term storage through persistent neural activity becomes potentially problematic when there is more than one to-be-remembered item^13^. Previous evidence has demonstrated that different to-be-remembered items can be simultaneously represented by *overlapping* neural populations^7–9,41^. Under these conditions, short-term storage through persistent neural activity could lead to representational conflicts. So how can overlapping neural populations simultaneously represent multiple to-be-remembered items, avoiding representational conflicts that might arise from persistent neural activity?

‘Activity-silent’ theories of working memory propose that the storage of to-be-remembered items can occur, in part, through short-term changes in synaptic weights^13–16,42,43^. Transient bursts of neural activity, rather than persistent neural activity, could serve to refresh these short-term synaptic changes. When overlapping neural populations are representing multiple to-be-remembered items, isolated bursts of neural activity—associated with different to-be-remembered items—could help to avoid representational conflicts. In support of these ideas, recent studies have confirmed that working memory delays are characterized by transient bursts of beta- (20-35 Hz) and gamma-band (45-100 Hz) activity in local field potentials^15,17,44^. Spiking activity associated with these transient bursts in beta- and gamma-band activity might play a role in refreshing short-term synaptic changes. It is important to note that models of working memory that incorporate such short-term synaptic changes (with transient bursts of neural activity)^42,45^ and models of working memory that incorporate persistent neural activity^40^ are not mutually exclusive. Both mechanisms could be contributing to the temporary storage of behaviorally important information^46,47^.

Here, we used high-density EEG (128 electrodes) and response times (RTs) to investigate whether rhythmic temporal coordination—like that previously observed during selective attention^3,4,31,33^—helps to organize transient neural activity associated with different to-be-remembered items. Rhythmic temporal coordination of neural activity would thereby help to avoid representational conflicts during working memory. Based on the notion that different item representations are refreshed at different oscillatory phases (and therefore separated in time)^18,48,49^, we specifically predicted that the relative strength of simultaneously held item representations would alternate over time as a function of oscillatory phase (Fig. 1D).

**Figure 1.**
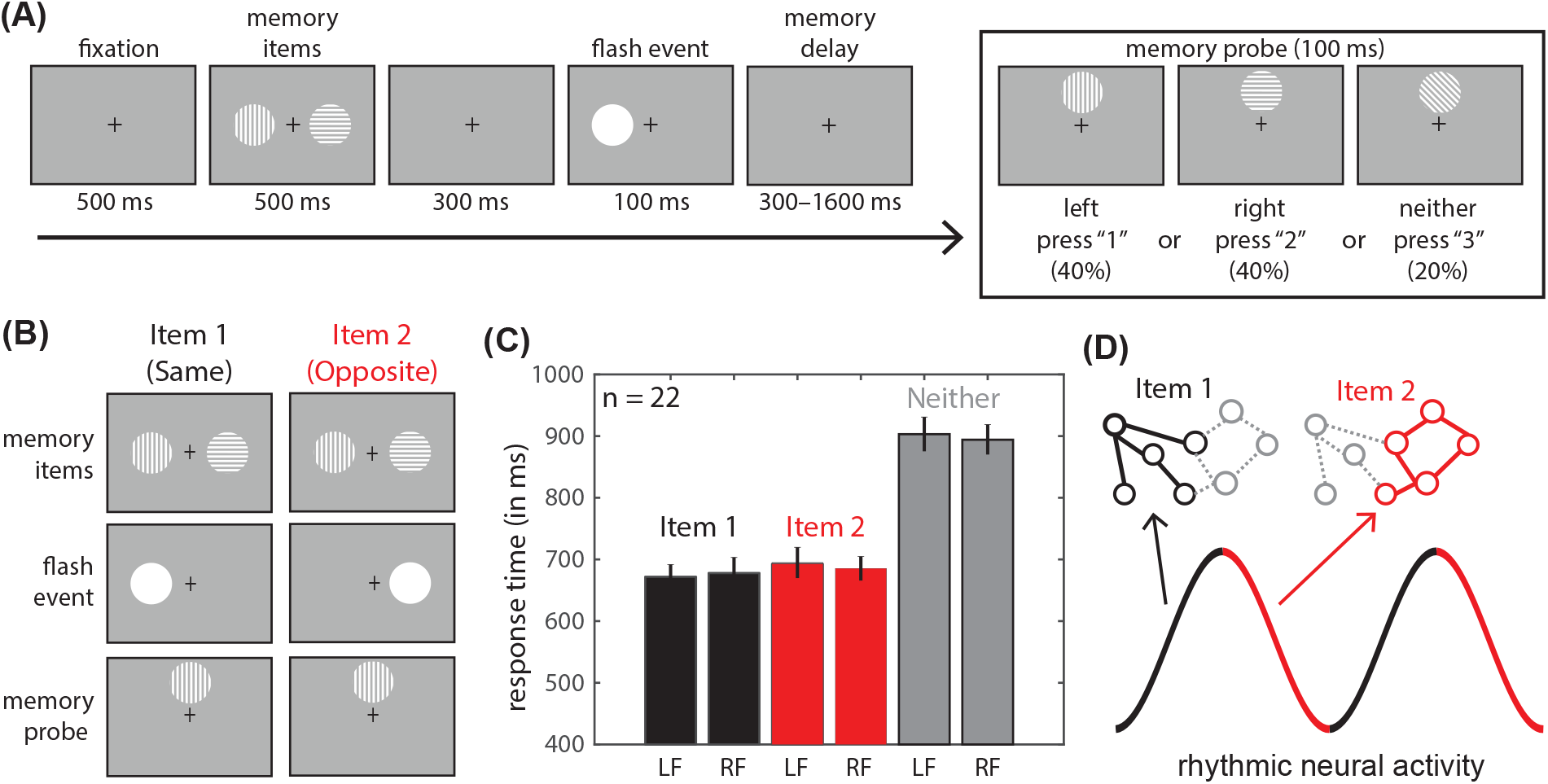
Behavioral performance during a working memory task. (A) Participants determined whether a probe was a match for either of two previously presented memory items. (B) We defined ‘Item 1’ trials as those trials when the probe matched the memory item presented on the same side of space as an intervening flash event and ‘Item 2’ trials as those trials when the probe matched the memory item presented on the opposite side of space as an intervening flash event. (C) Shows mean (n = 22) response times and the standard error of the mean for each condition (i.e., Item 1, Item 2, and Neither), depending on whether the flash event occurred to the left (LF) or right (RF) of central fixation. (D) We hypothesized that the relative strength of the representations for Item 1 (black) and Item 2 (red) would alternate in time as a function of oscillatory phase. In this schematic, circles represent cells and lines represent synaptic links within overlapping neural populations representing Item 1 (black) and Item 2 (red).

Consistent with this prediction, previous theoretical models of working memory have proposed that serially presented, to-be-remembered items are multiplexed at different phases of frequency-specific neural activity^18,48,49^. Supporting evidence for these theoretical models comes from both humans^20,21^ and monkeys^19^. For example, Bahramisharif *et al*.^20^ reported a link between item-specific, high-frequency band activity (75-120 Hz)—a proxy for population spiking^50^—and the phase of theta/alpha (7-13 Hz) oscillations. The specific phase associated with changes in item-specific, high-frequency band activity was dependent on when an item occurred within a sequence of to-be-remembered items. Recent behavioral studies in humans similarly reported that the internal representations of two objects in working memory are sampled and/or strengthened in alternating time windows, with this alternation occurring at a theta frequency (~6 Hz)^51–53^. While such behavioral findings suggest that the strength of internal representations fluctuate as a function of theta-band activity, the temporal binning typically used to measure human behavioral dynamics is too low to detect fluctuations at higher frequencies (i.e., > 15 Hz)^31,33,51–53^. Neurophysiological recordings in monkeys, which can investigate a much broader range of frequencies, have provided evidence that the strength of item representations fluctuates as a function of both lower and higher frequencies (i.e., 3 Hz and 32 Hz)^19^.

Here, we used an experimental task with simultaneously presented to-be-remembered items (Fig. 1A). The Research Subjects Review Board at the University of Rochester approved the study protocol. On each trial, differently oriented gratings (i.e., either horizontal, vertical, or diagonal) were presented 4 degrees from central fixation, on both the right and the left. Shortly after the presentation of these two memory items (i.e., to-be-remembered items), a task-irrelevant flash event was presented at the same location as one of the previously presented items (with equal probability). Here, we hypothesized that the relative strength of the item representations (i.e., neural representations of the to-be-remembered items) would alternate in time as a function of oscillatory phase (Fig. 1D). Flash events have previously been used to create consistent sampling patterns during attention tasks when there are multiple potential target locations^31^. The conceptualization of the flash event for the present task was also based on the use of retro cues during working memory tasks^54^. Retro cues have previously been used to boost the representation of one to-be-remembered item relative to other to-be-remembered items (during a memory delay). The flash event in the present task can be thought of as a retro cue that temporarily boosts the representation of one of the to-be-remembered items. As both items remain behaviorally relevant following the flash event (or retro cue), we hypothesized that the flash event would create a consistent pattern of alternation across trials, with the item on the same side as the flash event (i.e., the cued item) having a stronger initial representation than the item on the opposite side from the flash event (e.g., Item 1, Item 2, Item 1, Item 2…)^55^. Following a variable memory delay, participants (n = 22) reported, as quickly and as accurately as possible, whether a subsequent probe matched the memory item presented to (1) the left of fixation (40% of trials), (2) the right of fixation (40% of trials), or (3) neither of those memory items (20% of trials). The task therefore required that participants retain the spatial location and orientation of the two memory items (i.e., of the two visual gratings). To respond correctly, participants needed to sample internal representations of those previously presented, to-be-remembered items. The memory probe was presented at a neutral location, four degrees above central fixation. We defined ‘Item 1’ trials as those trials when the probe matched the memory item presented on the same side of fixation as the flash event, and ‘Item 2’ trials as those trials when the probe matched the memory item presented on the opposite side of fixation from the flash event (Fig. 1B).

Participants were able to perform the experimental task with high accuracy (mean = 85.2%, SE = 2.8). For the remaining analyses, we used RTs on correct trials as the behavioral measure (Fig. 1C). A two-way repeated measures ANOVA revealed a main effect for the item condition (i.e., Item 1, Item 2, or Neither, p < 0.001), no main effect for the flash condition (i.e., flash left or flash right, p = 0.31), and no interaction between the conditions (p = 0.68). Based on these results, we combined trials when the flash event occurred on either the right or left of central fixation. Follow-up t-tests revealed that RTs for both the Item 1 (p < 0.001) and the Item 2 (p < 0.001) conditions were significantly faster than RTs for the Neither condition, while RTs were not significantly different between the Item 1 and Item 2 conditions (p = 0.21). Previous work using visual search tasks has demonstrated that RTs are typically longer for ‘target absent’ trials than for ‘target present’ trials^56^, as more items must be searched, on average, during ‘target absent’ trials. Although the memory items in the present working memory task were not on the screen during the response period (Fig. 1A), the slower RTs for the Neither condition likely resulted, at least in part, from having to compare the memory probe with two non-matching item representations^57^. On Item 1 and Item 2 trials, the memory probe on half of the trials could be confirmed as a match after comparing it with only a single (and internal) item representation.

We next tested whether behavioral performance was linked to the phase of frequency-specific neural activity during the memory delay^29,58,59^, first combining all trials for which the memory probe was a match for either of the two memory items (i.e., combining the Item 1 and Item 2 conditions). We binned trials based on oscillatory phase just prior to when the memory probe was presented and then averaged z-standardized RTs (for each participant) within overlapping, 90-degree phase bins (with a step size of 10 degrees). Figure 2A shows RTs as a function of phase (i.e., a resulting phase-RT function) for a single electrode (i.e., D14, as labeled based on the BioSemi 128-channel ABC layout) and frequency, averaged across all participants (n = 22). We hypothesized that phase bins associated with peaks (i.e., slower RTs) and troughs (i.e., faster RTs) in behavioral performance would be separated by approximately 180 degrees. Based on this hypothesis, we fit the phase-RT functions with a one-cycle sine wave (Fig. 2A) and used the amplitude of that sine wave to measure the strength of the relationship between frequency-specific oscillatory phase and RTs^29,58,59^. Figure 2B shows the amplitude of these fitted sine waves (i.e., the strength of the phase-RT relationships) for all electrode (from 1-128) and frequency combinations (from 1-55 Hz), with statistically insignificant values set to zero. Supplemental Figure 1 (A and B) shows the same data without setting statistically insignificant values to zero. Statistical significance was based on a permutation test (see Methods). There were clusters of significant phase-RT relationships (p < 0.05, after correcting for multiple comparisons) in the theta (3-8 Hz) and beta bands (15-35 Hz), centered at 6 Hz and 25 Hz, respectively. Supplemental Figure 1 (C and D) more clearly demonstrates these peaks in the the theta and beta bands by averaging phase-RT relationships across all electrodes (n = 128). Theta- and beta-band activity have been previously linked to cognitive control^60,61^ and the maintenance^19,21,61–64^ of items held in working memory. Figure 2C shows scalp topographies for the electrodes with significant phase-RT relationships at 6 Hz and 25 Hz. Importantly, we also found consistent phase-RT relationships in the theta and beta bands when we subsequently split the data into the Item 1 and Item 2 conditions (Fig. 2D). Figure 2E shows all of the electrode and frequency combinations where we observed statistically significant phase-RT relationships for both the Item 1 and the Item 2 conditions, at either p < 0.05 or p < 0.1 (after corrections for multiple comparisons). The present results demonstrate that behavioral performance during a working memory task fluctuates over time as a function of oscillatory phase within multiple frequency bands. We propose that observed fluctuations in RTs reflect dynamic changes in the strength of the underlying item representations.

**Figure 2.**
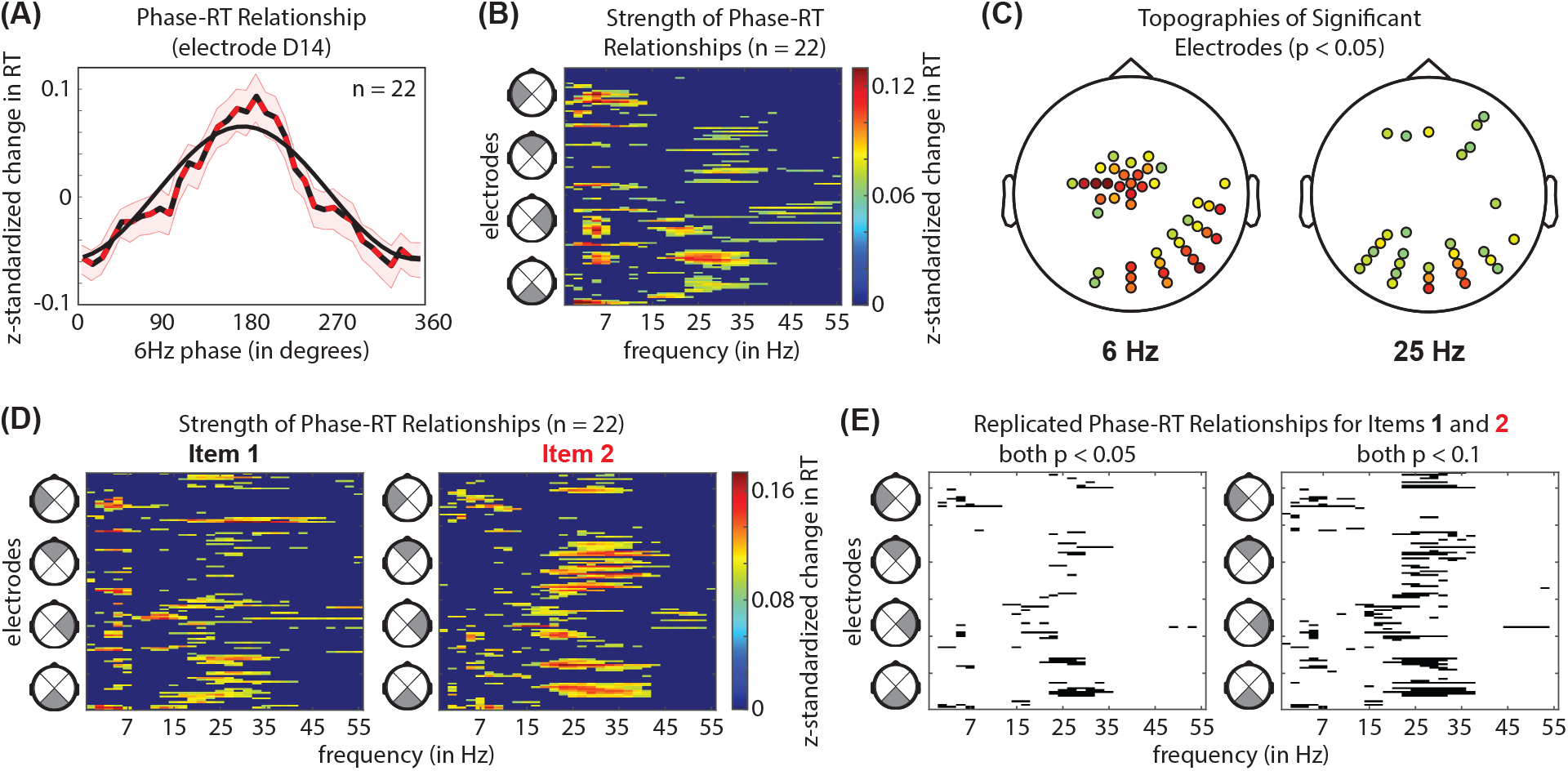
Response times (RTs) fluctuate as a function of theta and beta phase. (A) Illustrates the procedure for measuring the strength of phase-RT relationships, showing RTs as a function of oscillatory phase (at 6 Hz) for the electrode with the strongest phase-RT relationship (alternating red/black line). Here, we included all trials when the probe was a match for either of the two memory items (i.e., combining the Item 1 and Item 2 trials). The shaded region around the line represents the standard error of the mean. We used the amplitude of a fitted, one-cycle sine wave (solid black line) to measure the strength of the phase-RT relationship. (B) Shows the strength of statistically significant (p < 0.05) phase-RT relationships for all of the electrode (1-128) and frequency (3-55 Hz) combinations (statistically insignificant results were set to zero after corrections for multiple comparisons). There were clusters of significant results in both the theta and the beta bands, with (C) showing the scalp topographies of electrodes with significant phase-RT relationships within the theta (at 6Hz) and beta (at 25 Hz) bands. (B and C) Use the same color map to represent the strength of phase-RT relationships (i.e., as a z-standardized change in RT). (D) Results were consistent when calculated separately for the Item 1 and Item 2 conditions (statistically insignificant results were set to zero after corrections for multiple comparisons), with (E) showing overlapping, statistically significant results (in black) clustering in the theta and beta bands at p < 0.05 and p < 0.1 (after corrections for multiple comparisons).

We then used the item-specific phase-RT relationships to investigate whether the relative strength of the different item representations alternated in time as a function of oscillatory phase (Fig. 1D). Here, we tested whether the specific phases associated with better and worse behavioral performance were different for the Item 1 and Item 2 conditions (Fig. 3A). We again fit the participant-level phase-RT functions with one-cycle sine waves, but we now measured the phase of those sine-wave fits rather than the amplitude. We used a circular Watson-Williams test^65^ to determine whether the distributions of participant-level phases (i.e., the phase of the sine-wave fits) were different between the Item 1 and Item 2 conditions. We only tested for between-condition phase differences at electrode and frequency combinations where we previously detected significant phase-RT relationships for both the Item 1 and the Item 2 conditions (Fig. 2E). That is, we only made between-condition comparisons when both the Item 1 and the Item 2 conditions demonstrated a relationship between oscillatory phase and behavioral performance. Figure 3B shows the condition-specific phase distributions and angular means for a representative electrode and frequency. Figures 3C and 3D show all of the statistically significant (p < 0.05, after correction for multiple comparisons) electrode and frequency combinations, as well as the mean differences in phase between the Item 1 and Item 2 conditions. That is, the mean difference in the angular shift of the phase-RT functions between the Item 1 and Item 2 conditions. The analysis associated with the results shown in Figure 3C included all of the electrodes (e.g., 14 electrodes at 25 Hz) with significant phase-RT relationships at the p < 0.05 level for both the Item 1 and the Item 2 conditions (Fig. 2E), and the analysis associated with the results shown in Figure 3D included all of the electrodes (e.g., 33 electrodes at 25 Hz) with significant phase-RT relationships at the p < 0.1 level for both the Item 1 and the Item 2 conditions (Fig. 2E). The specific beta phases associated with better or worse behavioral performance were different for the Item 1 and Item 2 conditions. These findings are consistent with the hypothesis that the relative strength of different item representations alternates over time as a function of oscillatory phase (Fig. 1D). In comparison, we found no difference in the specific theta phases associated with better or worse behavioral performance between the Item 1 and Item 2 conditions (Fig. 3), despite earlier results demonstrating that RTs fluctuate over time as a function of theta phase (Fig. 2). Siegel *et al*.^19^ reported a similar pattern of results in the prefrontal cortex of monkeys, measuring information encoded by spiking activity as a function of oscillatory phase (rather than behavioral performance as a function of oscillatory phase). While information coding in spiking activity generally fluctuated as a function of both theta phase and beta phase, the optimal encoding of different items only varied as a function of beta phase. That is, there was no theta-band difference in the optimal encoding phase for different to-be-remembered items.

**Figure 3.**
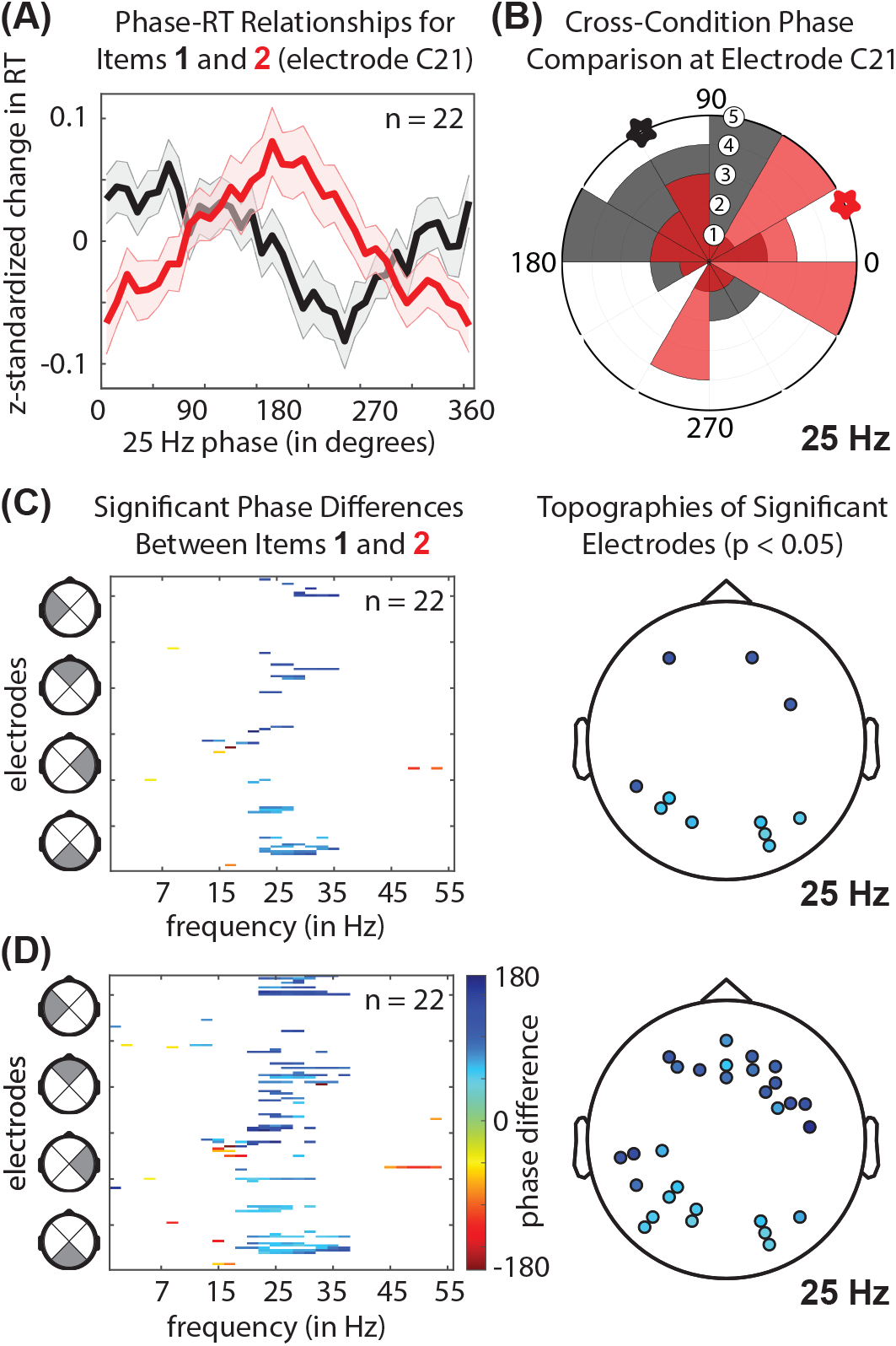
The beta phases associated with faster and slower response times (RTs) are different for the Item 1 and Item 2 conditions. (A) Shows RTs as a function of oscillatory phase (at 25 Hz), separately for the Item 1 (black line) and Item 2 (red line) conditions, for a single representative electrode. The shaded region around each line represents the standard error of the mean. (B) Shows circular histograms, plotting the phase of fitted one-cycle sine waves (see Fig. 2A) for each participant’s phase-RT functions (n =22), separately for Item 1 (in gray) and Item 2 (in light red). Overlapping measurements between the Item 1 and Item 2 conditions are represented in dark red. Participant counts in each phase bin ranged from 1 to 5. These results are shown for a single representative electrode, with the angular mean phases (n = 22) for each condition plotted as black (Item 1) and red (Item 2) stars. (C and D) Show all electrode (1-128) and frequency (3-55 Hz) combinations (in color) where the phases associated with faster and slower RTs were significantly different between the Item 1 and Item 2 conditions (p < 0.05 after corrections for multiple comparisons), including the scalp topographies of electrodes with significant effects in the beta band (at 25 Hz). (C and D) Use the same color map to represent the mean angular difference in phase between the Item 1 and Item 2 conditions. The analyses shown in (C) included all electrodes where we previously observed significant phase-RT relationships in both the Item 1 and the Item 2 conditions at the p < 0.05 level (see Figure 2E), while those shown in (D) included all electrodes where we previously observed significant phase-RT relationships in both the Item 1 and the Item 2 conditions at the p < 0.1 level (see Figure 2E).

A prominent model of oscillatory dynamics during working memory has proposed that theta-band activity coordinates item-specific higher-frequency activity^18,48^. According to this model, each cycle of the theta-dependent, higher frequency activity refreshes the representation of a different to-be-remembered item (see Bahramisharif *et al*.^20^ for supporting data). The number of cycles of the higher-frequency activity, nested within theta-band activity, might therefore reflect the number of to-be-remembered items. The present findings, as well as those from Siegel *et al*.^19^, instead indicate that each to-be-remembered item is refreshed within a *single* cycle of the higher frequency activity (i.e., within a single cycle of beta-band activity).

To test whether theta- and beta-band activity might be functionally linked in the present data, we measured whether beta amplitude varied as a function of theta phase (i.e., we measured phase-amplitude coupling^66,67^). Here, we used the same approach used to estimate phase-RT relationships. We calculated average amplitude (from 15-55 Hz) in overlapping theta-phase bins and fit the resulting phase-amplitude functions with one-cycle sine waves to measure the strength of the relationship (Fig. 4A)^29^. We specifically binned higher-frequency amplitude using theta phase (at 6 Hz) from the electrode (i.e., D14) where we previously measured the strongest phase-RT relationship (Fig. 2). Figure 4B shows the strength of phase-amplitude coupling (i.e., the amplitude of the sine-wave fits) for all of the statistically significant electrode and frequency combinations (p < 0.05 after corrections for multiple comparisons), with significant values clustered in the beta band. Figure 4C shows the scalp topography of the electrodes with significant coupling between theta phase (at 6Hz) and beta amplitude (at 25 Hz). These findings demonstrate that beta amplitude fluctuates as a function of theta phase. Such theta-dependent fluctuations in beta amplitude are consistent with transient changes in beta-band activity during a working memory delay, rather than a persistent increase in beta-band activity^15,17^.

**Figure 4.**
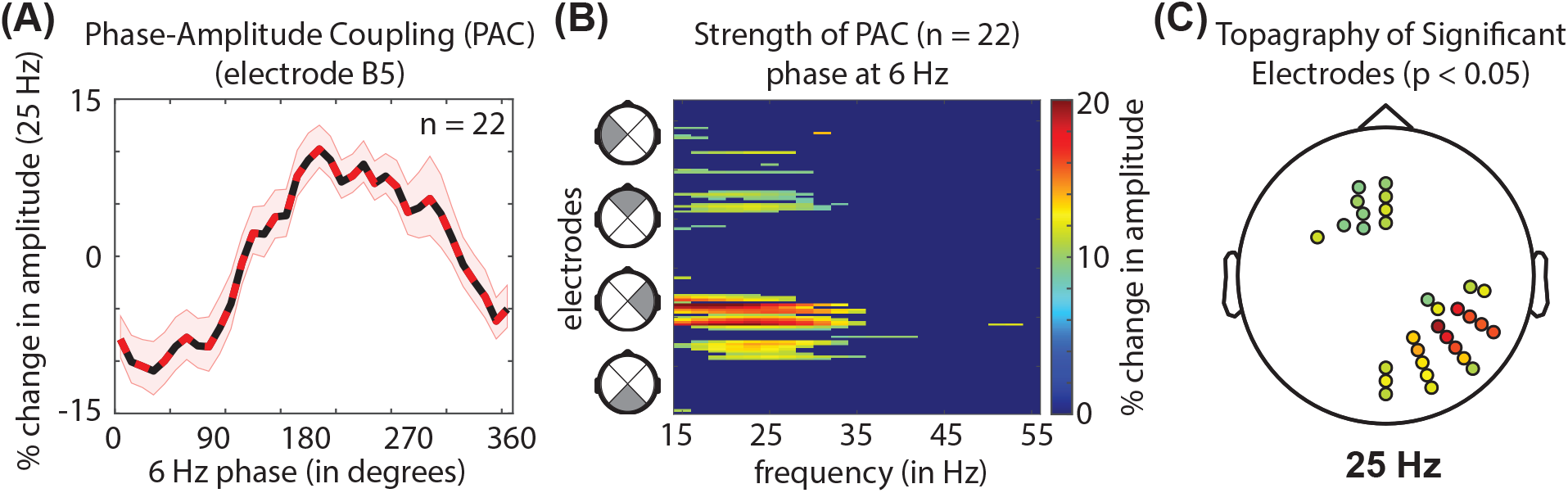
The amplitude of beta-band activity fluctuates as a function of theta phase. (A) Shows beta amplitude as a function of theta phase (at 6 Hz) for a single representative electrode (alternating red/black line), including all trials when the probe was a match for either of the two memory items (i.e., combining the Item 1 and Item 2 trials). The shaded region around each line represents the standard error of the mean. We used the amplitude of a fitted, one-cycle sine wave to measure the strength of the phase-amplitude relationship (see Fig. 2A). (B) Shows the strength of statistically significant (p < 0.05) phase-amplitude relationships for all of the electrode (1-128) and frequency (3-55 Hz) combinations. Statistically insignificant results were set to zero after corrections for multiple comparisons. For all phase-amplitude calculations, we used theta phase from the electrode that had the strongest phase-RT relationship (see Fig. 2A). There was a cluster of significant phase-amplitude results in the beta band, with (C) showing the scalp topography of electrodes with significant phase-amplitude relationships within the beta band (at 25 Hz). (B and C) Use the same color map to represent the percent change in beta-band amplitude (at 25 Hz) associated with different theta phases (at 6 Hz).

Figure 4A shows a representative phase-amplitude function from a single electrode, with beta amplitude peaking when theta phase is at approximately 180 degrees, which is the same theta phase associated with relatively worse behavioral performance (Fig. 2A). This pattern of results is potentially compatible with ‘activity-silent’ theories of working memory^13–16^. If theta-dependent increases in beta-band activity reflect transient neural activity that refreshes short-term synaptic changes, it would theoretically be most likely to occur when changes in synaptic weights are beginning to decay and behavioral performance is therefore at its worst (i.e., when item representations need to be refreshed). There are various mechanisms that contribute to short-term synaptic changes (e.g., calcium dynamics) on different timescales. Previous models of working memory have incorporated synaptic changes on the timescale of hundreds of milliseconds^42,45,46,68,69^, which might seem to be a mismatch for the frequency-specific modulations of behavioral performance reported here (Figs. 2, 3). The timescale of synaptic decay, however, can occur in stages, with a faster initial decay followed by a period of slower decay^64,65^. Moreover, refreshing events seem likely to happen on a shorter timescale than synaptic decay itself, helping to maintain and stabilize representations of to-be-remembered items before those representations fade. If theta-dependent beta-band activity is associated with neural events that refresh short-term synaptic changes, the minimum temporal separation for those refreshing events would be ~166 ms (i.e., the duration of a 6-Hz cycle). But the temporal separation of these refreshing events could be greater than 166 ms. While increased beta-band activity is associated with a specific theta phase (Fig. 4), it might not occur during every theta cycle. When increased beta-band activity does occur, we propose that it is associated with the sequential reactivation of multiple item representations (Fig. 3). Future work will need to investigate, for example, the specific cell types associated with these increases in beta-band activity.

The scalp topographies of our results are similar to previous EEG investigations of working memory. Supplemental Figure 2 shows the scalp topographies of isolated periodic components^70^ for the frequencies of interest during the memory delay (i.e., 6 Hz and 25 Hz). Theta-band activity in the present EEG results is consistent with frontal midline theta (FMT)^62^, which is thought to originate in medial prefrontal and anterior cingulate cortices^60,71^. FMT has been linked, for example, to memory capacity^61^ and behavioral performance^72,73^ during working memory tasks. Ratcliffe *et al*.^61^ recently provided evidence that FMT coordinates the maintenance of working memory content in posterior brain regions^38^, and neural synchronization is commonly observed between frontal and parietal cortices during working memory tasks^74–78^. Such neural synchronization can occur within both the theta and the beta bands^29,74–79^, consistent with the behaviorally relevant frequencies observed during the present working memory task (Figs. 2–4).

Theta-band activity within frontal and parietal cortices also coordinates rhythmically alternating states associated with either attentional sampling or shifting^28,29,31–34,80^, helping to temporally isolate sensory and motor functions of the attention network^3^. The present results provide evidence that the temporal coordination of neural activity helps to isolate item-specific neural activity during working memory. This is consistent with the rhythmic temporal coordination of neural activity being a more general mechanism for preventing conflicts during cognitive processes. Our findings show that behavioral performance during working memory, like that during selective attention^28,29,59^, fluctuates as a function of both theta phase and beta phase. Rhythmic temporal coordination of competing attentional states, however, seems to occur within the theta band (but not the beta band), whereas rhythmic temporal coordination of competing item representations seems to occur within the beta band (but not the theta band). Future research will be needed to test whether the behaviorally relevant, frequency-specific neural activity associated with selective attention (i.e., external sampling) and working memory (i.e., internal sampling) share common, frontoparietal neural sources. That is, whether there is a common clocking mechanism for organizing external and internal sampling. The theta phase associated with relatively worse working memory performance in the present task, for example, might reflect a theta-rhythmic switch in bias from internal to external sampling. In comparison, theta-dependent beta-band activity might reflect a process for maintaining short-term synaptic changes associated with working memory. It is still unclear whether and how these processes—the sampling of internal representations and the maintenance of internal representations—interact.

**Supplemental Figure 1.**
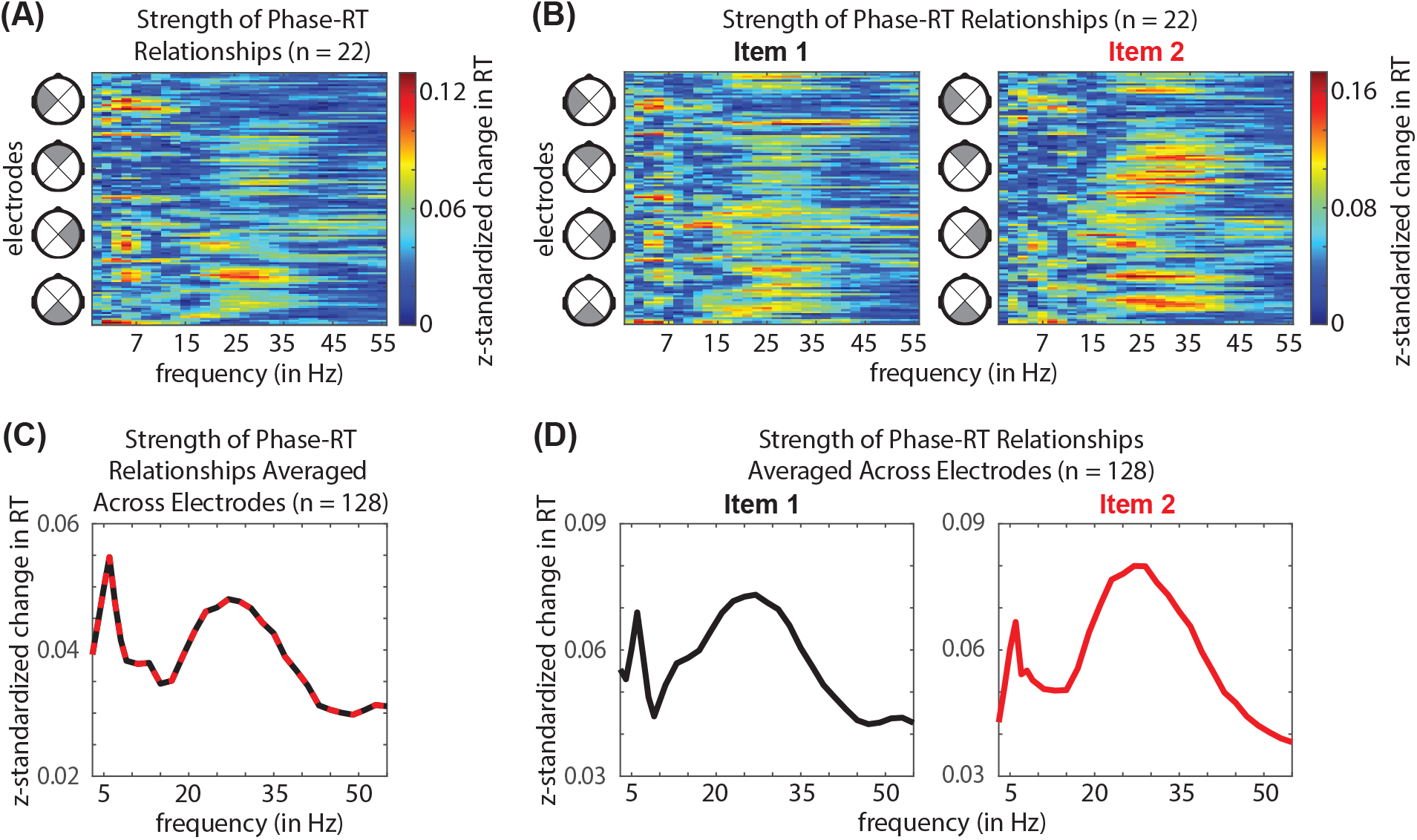
Response times (RTs) fluctuate as a function of theta and beta phase. (A) Shows the strength of phase-RT relationships (n = 22) for all of the electrode (1-128) and frequency (3-55 Hz) combinations. Here, we included all trials when the probe was a match for either of the two memory items (i.e., combining the Item 1 and Item 2 trials). (B) Results were consistent when calculated separately for the Item 1 and Item 2 conditions. (C and D) Show the results from (A and B) after averaging phase-RT relationships across all electrodes (n = 128). This additional averaging was included to more clearly illustrate the clustering of phase-RT relationships in the theta and beta bands.

**Supplemental Figure 2.**
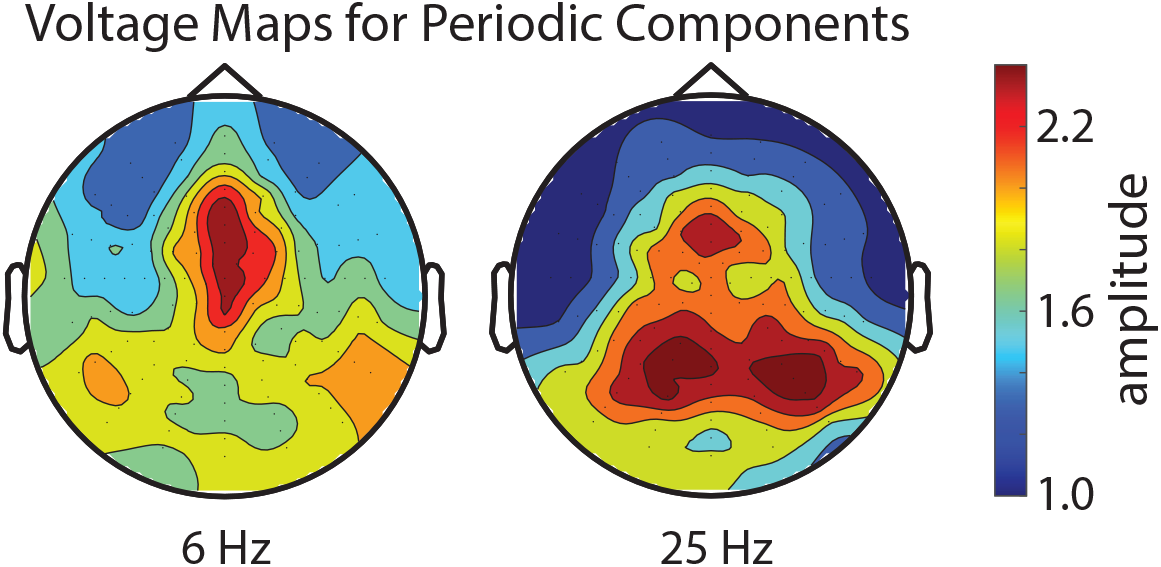
Scalp topographies of the isolated periodic components within the theta and beta bands. We used the FOOOF algorithm^70^ to remove the aperiodic, 1/f component, isolating the periodic components at behaviorally relevant frequencies. These scalp topographies are consistent with previous EEG studies that measured theta- and beta-band activity during a working memory delay.

## METHODS

### Contact for Resource Sharing

Further information and requests for code or data should be directed to and will be fulfilled by the Lead Contact, Ian C. Fiebelkorn.

### Subject Details

Twenty-seven individuals (12 females; 18-42 years old) with normal or corrected-to-normal vision and no history of neurological disease participated in the experiment. The Research Subjects Review Board at the University of Rochester approved the study protocol. Written informed consent was obtained from all participants prior to data collection, in line with the Declaration of Helsinki. Five participants were excluded from the analyses because of excessive blinks and/or eye movements (i.e., on greater than 20 percent of trials).

### Behavioral Task and Behavioral Data

Figure 1 summarizes our experimental design. The experiment was administered in a light- and sound-attenuated chamber. Presentation software (Neurobehavioral Systems, Albany, CA, USA) was used to control stimuli and monitor responses. The visual stimuli were presented on a 24-inch LCD monitor (ASUS Predator), operating at a refresh rate of 100 Hz. Participants began each trial by clicking the left mouse button. At the beginning of each trial a crosshair appeared at the center of the screen. Participants were instructed to maintain fixation throughout the duration of a trial and to try to blink between trials (i.e., withhold blinks during trials). After 500 ms, two memory items were presented (duration = 500 ms), one to the left and one to the right of central fixation (i.e., 4 degrees from central fixation). The two memory items, each with a diameter of 4 degrees, were differently oriented (i.e., horizontal, vertical, or diagonal) visual gratings (2.25 cycles per degree). A task-irrelevant flash event (duration = 100 ms) then occurred 300 ms after the presentation of the memory items, presented at the location of either of the previously presented memory items (with equal probability). Here, we hypothesized that the relative strength of the item representations (i.e., neural representations of the to-be-remembered items) would alternate in time as a function of oscillatory phase (Fig. 1D). We included the flash event to create a consistent pattern of alternation across trials (e.g., Item 1, Item 2, Item 1…)^31,81,82^. Following the flash event there was a variable memory delay, from 300-1600 ms. This variable memory delay was sampled from a uniform distribution. At the end of each trial, a memory probe (duration = 100 ms) was presented 4 degrees above central fixation. Participants reported, as quickly and as accurately as possible, whether the probe matched the memory item presented to (1) the left of fixation (40% of trials), (2) the right of fixation (40% of trials), or (3) neither of those memory items (20% of trials) by pressing 1, 2, or 3 on a keyboard. The participants were instructed to simultaneously place their fingers on the 1, 2, and 3 keys. An auditory “ding” was presented when participants responded correctly. We defined ‘Item 1’ trials as those trials when the probe matched the memory item presented on the same side of fixation as the flash event, and ‘Item 2’ trials as those trials when the probe matched the memory item presented on the opposite side of fixation from the flash event (Fig. 1B).

We tested whether response times (RTs) for correct trials differed depending on the Item condition (i.e., Item 1, Item 2, or Neither) and the Flash conditions (Right Flash or Left Flash) using a two-way repeated measures ANOVA and follow-up t tests. Based on the results of these statistical tests, we combined trials when the flash occurred on either the left or right of central fixation.

### Data Acquisition and Pre-processing

Electroencephalographic (EEG) data were acquired with a 128-channel ActiveTwo BioSemi system (Amsterdam, the Netherlands), sampling at a rate of 512 Hz. For all analyses, we used a combination of customized MATLAB functions (The MathWorks, Natick, MA, USA) and the Fieldtrip toolbox (Donders Institute for Brain, Cognition, and Behaviour, Radbound University Nijmegen, the Netherlands)^83^. The EEG data were first zero-padded (2.5 seconds) and epoched (from 2.6 seconds before the memory probe to 0.8 seconds after the memory probe), then linearly detrended and demeaned. A discrete Fourier Transform (DFT) filter was used to remove 60-Hz line noise, and the data were re-referenced to the average of all 128 electrodes (i.e., we used an average reference). The data were visually inspected for each subject to determine a voltage threshold for removing all trials with artifacts associated with eye movements or blinks, using electrodes positioned near the eyes. For trials without evidence of eye movements or blinks, a threshold of ± 100 μV was used to identify trials with other noise transients^58^. The data at individual electrodes were interpolated, using the nearest neighbor spline^84^, if fewer than 10% of electrodes were affected. Subjects who had artifacts on more than 20% of trials were excluded from all analyses (n = 5). The remaining subjects (n = 22) had an average of 8% of trials removed during artifact rejection, leaving an average of 880 trials per subject.

The present analyses were focused on frequency-specific phase during the memory delay (i.e., just prior to the memory probe). To measure phase on each trial, frequency-specific Morlet wavelets were used, with a varying number of cycles, from 2 cycles between 3 and 8 Hz, and increasing logarithmically from 2-5 cycles between 9 and 55 Hz. Each wavelet has a temporal extent based on the frequency and number of cycles. For example, a 4-Hz wavelet with 2 cycles extends for 500 ms. To limit the overlap between phase measurements and the evoked potential that occurred following the flash event, we only included trials where the memory delay was greater than 750 ms. For analyses measuring the relationship between frequency-specific phase and response times, the wavelet was fit for each frequency such that the last time point included in the phase measurement was the time point just prior to presentation of the memory probe. For the analysis of phase-amplitude coupling, wavelets for higher frequencies (i.e., 15-55 Hz) were centered at the same time point as the wavelet at 6 Hz (i.e., −167 ms). Pre-probe phase and amplitude measurements were calculated based on the complex output of the wavelet convolution (i.e., taking either the angle or the absolute value).

### Measuring Phase-Behavior and Phase-Amplitude Relationships

Here, we tested whether behavioral performance during a working memory task fluctuates as a function of oscillatory phase. To measure whether RTs were related to pre-probe phase (i.e., phase during the memory delay), RTs were first normalized for each participant by subtracting the mean of the RTs and dividing by the standard deviation of the RTs (i.e., we calculated z-standardized RTs). Here, trials with an RT less than 200 ms and trials with an RT greater than four standard deviations from the mean were excluded from further analysis^85^. For each condition (e.g., Item 1 and Item 2 combined, and Item 1 and Item 2 separately), RTs were binned based on frequency-specific pre-probe phase measurements (see the previous section for how we measured pre-probe phase), and average RTs were calculated in overlapping phase bins. The phase bins had a width of 90 degrees (e.g., 0-90 degrees) and were shifted forward in 10-degree steps (e.g., 10-100 degrees, then 20-110 degrees, etc.). This procedure was repeated to generate phase-RT functions, spanning all phases, for each frequency and each electrode. To capture consistent phase-RT relationships, these functions were averaged across participants (n = 22). Here, we predicted that the averaged phase-RT functions would have a signature shape, with a peak in RTs separated from a trough in RTs by approximately 180 degrees^29,58^. Based on this hypothesis, phase-RT functions were reduced to a single value for each frequency and electrode, using the following procedure: a discrete Fourier transform (DFT) was applied to each function (i.e., at each frequency and electrode) and the second component, representing a one-cycle sine wave (matching the hypothesized shape of the phase-detection relationship), was kept. The amplitude of this one-cycle, sinusoidal component—determined both by how closely the function approximated a one-cycle sine wave and by the effect size—was used to measure the strength of the phase-RT relationship^29,58^ (Fig. 2A).

An identical procedure was used to test for a relationship between theta phase (at 6 Hz) and higher-frequency amplitude (i.e., phase-amplitude coupling) during the memory delay (i.e., just prior to the memory probe). Here, phase-amplitude functions (rather than phase-RT functions) were generated by binning higher-frequency amplitude (from 15-55 Hz) by theta phase, then averaging higher-frequency amplitude within the overlapping theta-phase bins^29^. Here, higher-frequency amplitudes were exclusively binned based on pre-probe theta phase from the electrode where we previously measured the strongest phase-RT relationship (Fig. 2).

For both the phase-behavior and the phase-amplitude analyses, statistical significance was determined by iteratively shuffling (5000 times) the trial-level phase measurements (breaking the relationship between either phase and behavior or phase and amplitude). For each iteration, the analysis steps were then repeated, using shuffled data to calculate the strength of the phase-RT relationships or the phase-amplitude relationships. The resulting reference distributions (at each frequency and/or electrode) were compared to the magnitude of the observed data. For all analyses, before determining statistical significance, we controlled for the false discovery rate (accounting for multiple comparisons)^86^.

### Comparing Phase-RT Relationships Between Item Conditions

Here we predicted that the relative strength of the representations of different items should alternate in time as a function of oscillatory phase (Fig. 1D). We specifically tested whether the specific phases (from 3-55 Hz) associated with faster and slower RTs were different between the Item 1 and Item 2 conditions. Here, only electrodes where the phase-RT relationships exceeded a significance threshold of either p < 0.05 or p < 0.1 for both conditions (Fig. 2E) were included in the analysis. As described above, the discrete Fourier transform (DFT) was applied to each phase-RT function (see previous section) and the second component, representing a one-cycle sine wave, was kept. Here, the phase of the one-cycle sine waves (i.e., the angle of the second component), for each participant, were used (rather than the amplitudes). A circular Watson-Williams test^65^ determined whether the phases (n = 22) of the one-cycle sine waves were statistically different for the Item 1 and Item 2 conditions (Fig. 3B). That is, whether faster and slower RTs for the Item 1 condition were associated with different phases than faster and slower RTs for the Item 2 condition.

## ACKNOWLEDGMENTS

This work was supported by grants from the National Science Foundation (NSF 2120539) and the Searle Scholars Program to I.C.F. We would like to thank Dr. Edmund Lalor for inviting us to collect data in his laboratory while our own laboratory space was being renovated.

## AUTHOR CONTRIBUTIONS

I.C.F. conceived of the experiment. M.A. and Z.V.R. collected data. I.C.F analyzed the data. I.C.F wrote the first draft of the manuscript. I.C.F., M.A., and Z.V.R. edited the manuscript.

## DECLARATION OF INTERESTS

The authors declare no competing interests.

## Notes

### Competing Interest Statement

The authors have declared no competing interest.

### Summary of Updates

This version includes substantial revisions to the framing, interpretation, and presentation of the findings (i.e., the figures).

